# Domain-specific mechanisms of YAP1 variants in ocular coloboma revealed by in-vitro and organoid studies

**DOI:** 10.1101/2025.10.28.685193

**Authors:** Srishti Silvano, Annika Rick-Lenze, James Bagnall, Mrinalini Saravanakumar, Xinyu Yang, Robert Lea, Lindsay Birchall, Julie R Jones, Jessica M Davis, Anzy Miller, Rachel E Jennings, Elliot Stolerman, Jamie M. Ellingford, Simon C. Lovell, Forbes Manson, Gavin Arno, Panagiotis I. Sergouniotis, Cerys S Manning

## Abstract

The conserved transcriptional co-activator YAP1 is a central regulator of organ development and tissue homeostasis, integrating mechanical and biochemical cues to control cell proliferation and survival. YAP1 variants underlie a spectrum of congenital disorders, including autosomal dominant coloboma that can occur alone or with syndromic features. Despite this clinical significance, the functional role of YAP1 in human eye development, as well as the impact of disease-associated missense variants, remains poorly understood. Here we show YAP1 expression at the optic fissure in human embryos, a key structure involved in coloboma pathogenesis. We also identify a novel YAP1 variant in a proband with syndromic coloboma and investigate five previously reported coloboma-associated YAP1 variants. Using in silico prediction, cell-based assays, and fluorescence cross-correlation spectroscopy (FCCS) to directly quantify YAP1–TEAD binding, we demonstrate that the position of YAP1 missense variants dictates their functional changes. TEAD-binding domain mutations most strongly disrupted transcriptional activity in a luciferase assay, whereas all tested variants impaired induction of endogenous YAP1-TEAD target genes. Furthermore, mimicking reduced YAP1-TEAD binding using verteporfin small molecule in retinal organoids led to reduced progenitor proliferation and survival. These findings establish defective YAP1-dependent transcription as a mechanism driving congenital eye malformations and provide a framework for interpreting the pathogenicity of human YAP1 variants. More broadly, this study highlights the need for functional analyses to connect genetic variation with disease.

## 1. INTRODUCTION

Ocular coloboma is a rare congenital eye malformation with an estimated prevalence of around 1 in 5000 births [1]. It often leads to significant visual impairment, making patients’ everyday life challenging and contributing to approximately 10% of childhood blindness [2]. Coloboma arises from incomplete closure of the optic fissure during embryonic development, resulting in a gap in ocular structures such as the iris, choroid, retina, or optic nerve. Furthermore, co-occurence of coloboma with other tissue fusion defects: cleft lip and/or palate (CL/P) is often reported [3, 4]. Coloboma has been linked to pathogenic variants in more than 40 genes involved in eye development [5, 6]. *YAP1* (Yes-associated protein 1), a transcriptional co-activator in the Hippo signalling pathway, plays a key role in eye development in animal models [7, 8], and variants in *YAP1* have been associated with autosomal dominant coloboma in human [4, 9–11].

YAP1 regulates key processes such as cell proliferation, differentiation, and survival during embryonic development and is also shown to be a regulator of epithelial-mesenchymal transition (EMT) [9, 12, 13]. It’s subcellular localisation is dynamically regulated by phosphorylation, primarily through the Hippo kinase cascade [14–16]. When activated, the Hippo pathway kinases LATS1/2 bind to YAP1’s WW domains and phosphorylate conserved N-terminal serine residues, particularly Ser127 [17]. This phosphorylation triggers 14-3-3 protein binding, sequestering YAP1 in the cytoplasm and promoting its degradation [18]. Conversely, when LATS1/2 are inactive, YAP1 accumulates in the nucleus, where it binds transcription factors TEAD1-4 via its TEAD-binding domain. This upregulates key target genes such as *CTGF*, *AXL*, *CYR61*, and anti-apoptotic factors from the *Bcl2* and *AIP* families [15, 16, 19]. YAP1’s localisation is further modulated by physical cues such as substrate stiffness, cell area, and cell density [20]. This is in part mediated by interactions with membrane-associated proteins such as α/β-catenins and angiomotin (AMOT) family proteins which can promote YAP1 localisation at tight junctions [20]. These regulatory mechanisms ensure that YAP1 integrates biochemical and biomechanical signals to control cell fate.

YAP1 plays essential roles in eye development, regulating optic cup morphogenesis, retinal pigment epithelium (RPE) differentiation, and retinal progenitor expansion. In zebrafish, the loss of Yap1 and the closely related transcription factor Taz leads to coloboma and complete absence of the RPE [7]. Furthermore in mouse, conditional knockout of YAP1 causes a more severe phenotype of coloboma and microphthalmia. In the developing optic vesicle, a loss of pigmentation, similar to zebrafish, together with transdifferentiation of presumptive RPE cells into neural retina was observed [8]. Furthermore, YAP1-deficient retinal progenitor cells showed disrupted apical cell junctions, reduced proliferation and increased cell death [8]. To date, the role of YAP1 has not been systematically investigated in human retinal organoids (ROs), providing an opportunity to clarify its functions in early human retinal development, complementing findings from animal models.

Genetic studies have identified heterozygous YAP1 variants in multiple individuals with coloboma. These variants often show incomplete penetrance, where some carriers have no eye defects, and variable expressivity, meaning that the severity and associated features can differ widely between affected individuals. Williamson *et al*. identified co-segregating heterozygous nonsense mutations in two families, one with isolated coloboma and another with coloboma and extraocular features [4]. Also, DeYoung *et al*. reported a YAP1 frameshift variant in a proband with bilateral coloboma accompanied with microphthalmia [21].

Furthermore, Holt *et al*. reported a heterozygous single base-pair deletion in a proband with coloboma (inherited from his unaffected father) [11]. Similarly, Oatts *et al*. reported a heterozygous missense YAP1 variant in two maternal half-brothers affected by coloboma. The variant was inherited by their unaffected mother [10]. Additional disease-implicated *YAP1* variants have been deposited to clinical databases like ClinVar. The variable penetrance of *YAP1* variants in coloboma complicates determining whether a given variant is truly disease-causing or benign. Notably, despite recent advances in in-silico prediction tools, assessing the pathogenicity of missense variants remains particularly challenging. Such variants can disrupt protein function through diverse mechanisms, including altered folding, reduced stability, aberrant subcellular localisation, or impaired biochemical activity. Functional testing is therefore key to improving the accuracy of missense variant classification.

Here, we test a set of missense YAP1 variants associated with coloboma in cell culture assays and mimic the most severe TEAD binding domain variants in early-stage YAP1 retinal organoids. We find that inhibiting YAP1-TEAD binding in organoids significantly impacts the proliferation and survival of retinal progenitors. Additionally, we show that variants in the TEAD binding domain exert the most pronounced effects on YAP1 transcriptional activity. These findings highlight the pivotal role of the YAP1–TEAD interaction in human retinal development, providing a framework for understanding the pathogenicity of disease-associated YAP1 variants.

## 2. MATERIALS AND METHODS

### 2.1 Variant selection

Studied variants were identified through a ‘YAP1’ search in Clinvar (accessed 01/03/2023) with filtering for missense changes and further selection of variants in individuals with an eye anomaly. In addition, a published coloboma-associated YAP1 missense variant was also included (Table 1)[10].

**Table 1:** YAP1 variants shortlisted for this study.

| Protein change | mRNA change | dbSNP ID | gnomAD variant ID | gnomAD frequency | YAP1 domain | Isoform affected | Clinical phenotype | ClinVar interpretation | Reference |
| --- | --- | --- | --- | --- | --- | --- | --- | --- | --- |
| Valine 55 Leucine (p. Val55Leu) | c.163G>T | rs1190867473 | 11-102111011-G-T | 1/1599450 | TEAD binding-site | all | Uveal coloboma-cleft lip and palate-intellectual disability (COB1) | VUS | ClinVar |
| Methionine 86 Threonine (p. Met86Thr) | c.257T>C | rs2135086778 | - | - | TEAD binding-site | all | COB1 | Pathogenic | ClinVar |
| Phenylalanine 95 Serine (p. Phe95Ser) | c.284T>C | - | - | - | TEAD binding-site | all | Coloboma | N/A | Oatts et al. 2017 |
| Lysine 254 Arginine (p. Lys254Arg) | c.761A>G | rs1947930460 | - | - | WW2 domain | YAP1-2α(208), YAP1-2β(209), YAP1-2γ(201), YAP1-2δ(210) | COB1 | VUS | ClinVar |
| Serine 257 Phenylalanine (p. Ser257Phe) | c.770C>T | rs1947931415 | - | - | WW2 domain | YAP1-2α(208), YAP1-2β(209), YAP1-2γ(201), YAP1-2δ(210) | COB1 | VUS | ClinVar |
| Glutamic acid 356 Aspartic acid (p.Glu356Asp) | c.1068G>T | rs1950059078 | 11-102223657-G-T | 13/1614156 | Transcriptional activation domain | all | Coloboma, cleft lip and alveolar ridge, developmental delay | N/A | Novel variant |

COB1: uveal coloboma-cleft lip and palate-intellectual disability; VUS: variant of unknown significance

### 2.2 Exome sequencing for patient variant identification

The Agilent SureSelect^XT^ Clinical Research Exome (Human V5) kit was used to target known disease-associated exonic regions of the genome (coding sequences and splice junctions of known protein-coding genes associated with disease, as well as an exomic backbone) using genomic DNA from the submitted sample. Sequencing was performed on the Illumina NextSeq500 instrument. Using NextGENe^®^ software and an in-house bioinformatics pipeline, the DNA sequence was aligned and compared to the human genome build 19 (hg19/NCBI build 37). The Cartagenia Bench Lab NGS software (now known as Alissa Interpret) was used to filter and analyse sequence variants identified in the patient. The reported variants were confirmed by Sanger sequencing for the patient and her mother.

### 2.3 Computational pathogenicity prediction and protein modelling of YAP1 variants

A set of *in-silico* prediction tools was used to classify YAP1 variants. These included: CADD [22], REVEL [23], SIFT [24], Mutation Taster [25], AlphaMissense [26]. Protein conservation scores were also used for the variant classification. The scores were available at dbNSFP, ProtVar UI V1.3, and VarSome [27–29].

To visualise the impact of the studied genetic variants, we used YAP1 structure models 3KYS (YAP1-TEAD complex) and 5YDY (YAP1-LATS complex), both extracted from the Protein Databank (PDB)[30, 31]. Hydrogen atoms were added with the ‘Reduce’ software [32] and interactions between the variant amino acid and the rest of the protein were visualised with Probe [33]. In each case, the side chain conformation was modelled with a rotamer that gave the most favourable interactions [34].

### 2.4 YAP1 site-directed mutagenesis

Plasmids pLL3.7-EF-EYFP-YAP1_WT-PolyA (Addgene, #112284)(hereby EYFP-YAP1-WT) and pLL3.7-EF-EYFP-YAP1_S94A-PolyA (Addgene, #112286)(YAP1 p.Ser94Ala) were gifts from Erik Sahai [35]. The following YAP1 variants p.Val55Leu (c.163G>T), p.Met86Thr (c.257T>C), p.Phe95Ser (c.284T>C), p.Ser127Ala (c.379T>G), p.Lys254Arg (c.761A>G), p.Ser257Phe (c.770C>T), p.Glu356Asp (c.1068G>T) were introduced to the wildtype plasmid EYFP-YAP1-WT using Q5® Site-Directed Mutagenesis Kit (New England Biolabs, #E0554). All plasmids were sequence-verified using Sanger sequencing (Eurofins Scientific) and whole plasmid Nanopore sequencing (Source BioScience).

### 2.5 Cell culture maintenance and transfection for Luciferase assay

HEK293 LTV cell line (Cell Biolabs, #LTV-100) was maintained in DMEM high glucose (Sigma-Aldrich, #D6429) with 10% foetal bovine serum (FBS; Gibco, #10500064) and 1 % penicillin/streptomycin (Merck Life Sciences, #P4333). For the TEAD-dependent luciferase assay, HEK293 LTV cells at 70-80% confluency were plated on fibronectin (10μg/ml; Sigma-Aldrich, #341635) coated 12-well plates (Corning, #3513) at a density of 60,000 cells/well for 48 hours. These were then transfected with 100 ng 8xGTIIC-luciferase (Addgene, #34615) and 50ng pRL-CMV-*Renilla* luciferase control vector (Promega, #E2261) with 150ng of EYFP-YAP1 constructs (wildtype, p.Val55Leu, p.Met86Thr, p.Ser94Ala, p.Phe95Ser, p.Ser127Ala, p.Lys254Arg, p.Ser257Phe, p.Glu356Asp) in duplicates using the Lipofectamine^®^ 3000 transfection kit (Invitrogen, #L3000-008). The cells were cultured for 24 hours, then lysed with 200ul 1X passive lysis buffer (Promega, #E1941); 20 μl lysate was transferred to luminometer-compatible Nunc^TM^ microwell white polystyrene 96-well plate (Thermofisher Scientific, #236105) to perform Luciferase assay using the kit Dual-Glo Luciferase assay system (Promega, #E2920). The luciferase luminescence was normalised with *Renilla* luminescence. GloMax^®^ navigator microplate luminometer (Promega, #GM2010) was used to measure the luminescence. Each variant was assayed three times in duplicates.

### 2.6 Lentivirus preparation and transduction

EYFP-YAP1 wildtype and variant plasmids were transfected into HEK293 LTV cells upon reaching 70% confluency in a 10cm cell culture plate following lentivirus safety protocol. Briefly, 1X polyethylenimine (PEI STAR^TM^; R&D systems, #7854) was mixed with DMEM and incubated for 2 mins at room temperature. In a separate tube, DMEM was mixed with second-generation packaging vector MD2.G (6 μg)(Addgene, #12259) and VSV-G envelop expressing plasmid psPax2 (8μg)(Addgene, #12260), and EYFP-YAP1 construct and incubated for 2 mins at room temperature. This was followed by mixing the solutions from both tubes together (vector-PEI) and incubating for 30 mins at room temperature. The HEK293 LTV cell media was replaced with half of the regular volume and cells were transfected with the vector-PEI solution dropwise and incubated overnight at 37 °C; 5% CO2. Next day, the cell culture media was replaced with the full volume of media containing 1M sodium butyrate (100ul in 10ml cell culture media)(Sigma-Aldrich, #303410) and incubated for 6 hrs. The cell culture media was subsequently replaced with fresh complete media and incubated for 2 nights. The cell culture media was then filtered through a 0.45um syringe filter (Starlab, #E4780-1453) in a tube and virus was concentrated with 1X polyethylene glycol (PEG-it^TM^; SBI System Biosciences, #LV810A-1) for 4 days at 4°C. Concentrated viral supernatant was used to transduce HEK293 LTV cells for 48 hrs., which were subsequently FACS sorted to enrich for EYFP positive cells.

### 2.7 Protein separation and Western blotting

HEK293 LTV cells transfected with EYFP-YAP1 constructs were subjected to lysis with Pierce RIPA lysis buffer (Thermofisher Scientific, #89900) with protease inhibitor (Roche, #11836170001). The supernatant from cell lysates was mixed with NuPAGE LDS sample buffer (Invitrogen, #NP0007) with NuPAGE sample reducing agent (Invitrogen, #NP0004) and incubated at 95^0^C for 5 mins. The protein samples were loaded in Mini-PROTEAN TGX stain-free gels (Bio-Rad, #4568125) in trisglycine/SDS buffer (BioRad, #1610772) and electroporation was run at 100 V for 1.5 hr. The protein bands were transferred to a nitrocellulose membrane (BioRad, #1704158) in TransBlot turbo transfer system. The blot was blocked with 5% milk in 1X Tris buffered saline (BioRad, #1706435)-Tween 20 (Sigma Life Science, #P9416). Antibodies used were rabbit anti-YAP (D8H1X)(1:1000; Cell Signaling, #14074), mouse anti-β-actin (1:1000; Sigma-Aldrich, #A1978), anti-mouse IgG HRP-linked (1:2000; Cell Signaling, #7076) and anti-rabbit IgG HRP-linked (1:2000; Cell Signaling, #7074). The blot was visualised with Pierce^TM^ ECL western blotting substrate (Thermofisher Scientific, #32209) in the ChemiDoc MP imaging system (BioRad).

### 2.8 Fluorescence cross-correlation spectroscopy (FCCS)

Fluorescence cross-correlation spectroscopy (FCCS) images were obtained using the ZEN 2.1 software on a Zeiss LSM 880, equipped with a stage-mounted incubator to maintain cells at 37 °C in a humidified 5% CO2 atmosphere.-For FCCS, FACS-sorted HEK293 LTV cells with EYFP-YAP1 wildtype and variant constructs were plated at 60,000 cells/well of a fibronectin (10μg/ml) -coated 12-well plate. The cells were then transfected with 150ng or 225ng of TEAD1-mscarlet-I plasmid for 48 hours. The cells were then plated at 35,000 cells/well on fibronectin-coated 4-chambered glass-bottom plate (Greiner bio-one, #627870) for 48 hours, which was followed by FCCS imaging after a medium change. FCCS measurements were obtained as previously reported [36]. In brief, points were measured using Zeiss inverted LSM880 confocal microscope equipped with a 40x 1.3N.A. oil immersion objective with a pinhole set to 1 airy unit. The fluorophores, EYFP and mScarlet-I, were excited using a 488nm and a 561nm laser line respectively. Care was taken to use appropriate power to reduce bleaching whilst maintaining signal. FCCS measurements were performed in each cell nucleus using acquisition times of 10 secs for 1 repetition and a collection volume of 1 airy unit calibrated in the x-y plane for maximum signal intensity. Data were analysed using a previously reported python script to fit correlation curves generated by Zen acquisition software [37].

### 2.9 Fluorescence imaging for YAP1 variants nuclear vs cytoplasmic localisation ratio

HEK293 LTV cells with EYFP-YAP1 wildtype and variant constructs were plated at 45,000 cells/well on fibronectin (10μg/ml) coated 4-chambered glass-bottom plate for 48 hours. The cells were then fixed and counter-stained with DAPI (Sigma Aldrich, #D9542) to image using Zeiss LSM880. Fluorescence intensity was measured using FIJI software. ROIs were selected for nucleus and cytoplasm for each cell and all measures were background-corrected using fluorescence intensity outside of the cell area.

### 2.10 Human iPSCs maintenance and differentiation to retinal organoids

The human induced pluripotent stem cell (hiPSC) wild-type AD4 line was a gift from Majlinda Lako, Newcastle University, UK. Cells were grown on Vitronectin (Thermofisher Scientific, #A14700) coated dishes and maintained in StemFlex™ (Thermofisher Scientific, #A3349401) at 37°C and 5% CO2. Before the confluence reached beyond 70%, hiPSCs were passaged at a ratio of 1:6 in StemFlex™ with 10uM Y-27632 ROCK inhibitor (ROCKi; Sigma, #Y0503) using 0.5mM EDTA. ROs were generated according to the method of Chichagova *et al*. with few modifications [38]. On day -2, hiPSCs were dissociated into single cells using accutase (Sigma, #A6964) and seeded at a density of 7000 cells/well of a 96-well Nunclon Sphera-Treated U-bottom plate (Thermofisher Scientific, #174925) in StemFlex™ with 10uM ROCKi. Two days later, StemFlex™ was replaced with 200μl of differentiation medium and renewed every second day. Differentiation medium consists of 41% Iscove’s modified Dulbecco’s medium (Gibco, #12440-046), 41% Hams F-12 (Gibco, #11765-047), 15% KnockOut serum replacement (Gibco, #10828010), 1% GlutaMax (Gibco, #35050061), 1% chemically defined lipid concentrate (Gibco, #11905031), 1% pen-strep, and 225μM 1-Thioglycerol (Sigma, #88640). At day 6, differentiation medium was supplemented with 50ng/ml BMP4 (R&D Systems, #314-BP-010). Medium was then renewed every third day. On day 18 of differentiation, medium was substituted with maintenance medium containing DMEM/F-12 (Thermofisher, #31330095), 5% FBS, 1% N2 supplement (Thermofisher Scientific, #17502048), 0.1 mM taurine (Sigma, #T8691), 0.25μM retinoic acid (Sigma, #R2625), 0.25μg/ml Fungizone (Gibco, #15290018), 1% pen-strep. Medium was renewed three times per week. For verteporfin treatment, day 35 ROs were treated for seven days with 0.25µM verteprofin (Merck, #SML0534), or for the control group with an equivalent volume of DMSO. For Selinexor (SLX) treatment, day 35 ROs were treated for 24hrs with 1µM SLX, or for the control group with an equivalent volume of DMSO.

### 2.11 Immunostaining of retinal organoids and human embryonic tissue sections

ROs were collected and fixed in 4% paraformaldehyde for 20 mins while rocking at room temperature, followed by incubation in 30% sucrose at 4°C overnight. ROs were then embedded in moulds with OCT Embedding Matrix (Pioneer Research Chemicals Ltd) and frozen at -80°C. Cryosections were cut at 12µm using Leica CM3050 S cryostat.

Human embryonic material was collected from medical and surgical terminations of pregnancy under ethical approval from the North West Research Ethics Committee (23/NW/0039) and under the codes of practice issued by the Human Tissue Authority and legislation of the UK Human Tissue Act 2008. Appropriate informed consent was obtained. Specimens were fixed within 1hr in 4% paraformaldehyde before processing and embedding in paraffin wax. Sectioning took place at 5μm intervals, with every eighth section taken for hematoxylin and eosin staining to confirm morphology and anatomical landmarks. Sections containing ocular tissue at CS15-16 were chosen for immunostaining. The sections were rehydrated in xylene and ethanol, and then, for antigen retrieval, were boiled in 1X sodium citrate for 10 mins and removed from heat for 20 mins.

Primary antibodies used on ROs were mouse anti-VSX2 (Santa Cruz Biotechnology, #sc-365519), rabbit anti-YAP1 (D8H1X)(1:200; Cell Signaling, #14074), mouse anti-Phospho histone H3 (1:200; Abcam, #Ab14955) and rabbit anti-cleaved Caspase3 (1:100; Cell Signaling, #9661), and on human tissue were mouse anti-YAP (63.7)(1:100; Santa Cruz Biotechnology, #sc-101199) and rabbit anti-p-YAP (Ser127)(1:1600; Cell signaling, #D9W2I). The secondary antibodies used were donkey anti-rabbit 488 (1:800; Invitrogen, #A21206) and donkey anti-mouse 594 (1:800; Invitrogen, #A21203). ROs were counterstained with DAPI, and the human embryonic tissues were dehydrated and mounted with Vectashield antifade mounting medium with DAPI (Vector Laboratories, #H1200).

### 2.12 Retinal organoid RNAseq and analysis

At each timepoint, eight ROs were collected and total RNA extraction was carried out using the QIAGEN® RNeasy Micro Kit, following the manufacturer’s protocol (Qiagen, #74104). Quality and integrity of the RNA samples were assessed using a 4200 TapeStation (Agilent Technologies) and Qubit (Thermofisher Scientific). Sequencing libraries were generated using the *Illumina*^®^ *Stranded mRNA Prep. Ligation* kit (Illumina) according to the manufacturer’s protocol by Genomic Technologies Core Facility at The University of Manchester. Paired-end sequencing (2× 75bp) was then performed on an Illumina NovaSeq6000 instrument. The output data was demultiplexed and BCL-to-Fastq conversion was performed using Illumina’s *bcl2fastq* software, version 2.20.0.422. The Genotype-Tissue Expression (GTEx) analysis pipeline (GTEx Consortium, 2019) was followed for quality control of the reads, alignment and expression quantification [39]. Briefly, alignment of the human reference genome GRCh38 was performed using STAR v2.7.4a [40]. The GENCODE v38 annotation was utilised for gene-level expression quantification [41]. RSEM v1.3.0 was used for transcript-level quantifications (in transcripts per million, TPM) [42].

### 2.13 Statistics

Statistical testing was performed in GraphPad Prism 9.3.1. Data was tested for normality using the Shapiro-Wilk method before selecting the appropriate test. Significance was defined as ns p>0.05, ^∗^p<0.05, ^∗∗^p<0.01, ^∗∗∗^p<0.001, ^∗∗∗∗^p<0.0001. Outliers were removed in the subcellular localisation assay using ROUT.

## 3. RESULTS

### 3.1 YAP1 is expressed in human retinal progenitor cells and RPE at the optic fissure

Given that coloboma arises from defects in optic fissure closure, we analysed YAP1 expression in human embryonic eye tissue during this developmental stage (CS15-16). We found that YAP1 was expressed throughout the presumptive neural retina and RPE. There was enrichment at the apical surface of the retina and hinge-region between the neural retina and the RPE (Fig.1A). In the developing neural retina, YAP1 qualitatively displayed predominantly cytoplasmic localisation. This was supported by the presence of phosphorylated YAP1 at Ser127 (phospho-YAP1), a marker of cytoplasmic YAP1 sequestration and active Hippo signalling (Fig. 1A). In contrast, the periocular mesenchyme qualitatively showed higher levels of total YAP1 relative to phospho-YAP1, suggesting a greater proportion of active, nuclear-localised YAP1 in this region. Meanwhile, developing lens tissue exhibited more cytoplasmic/transcriptionally inactive phospho-YAP1 (Fig. 1A).

**Figure 1.**
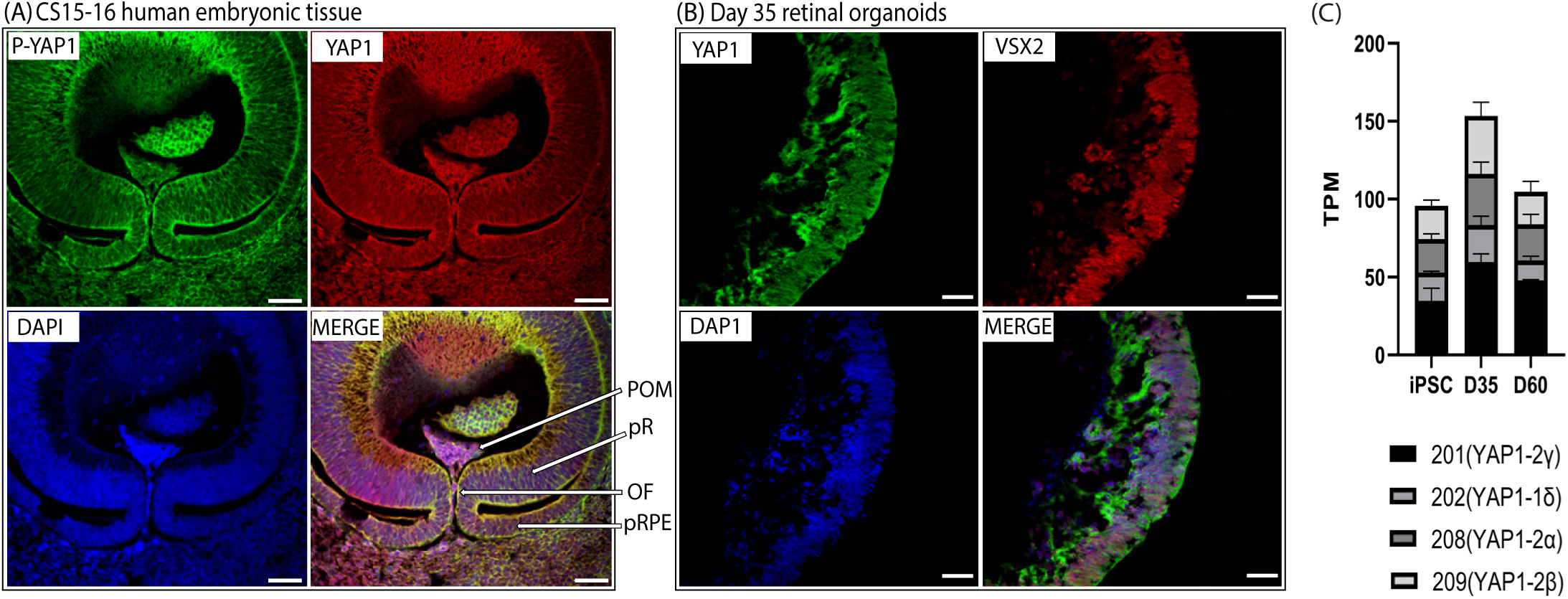
YAP1 expression during early human eye development. (A) Expression of phospho-YAP1 (P-YAP1(Ser127)) and YAP1 in human embryonic ocular tissue (CS15-16)[Scale: 50µm]. (B) Expression of YAP1 co-localised with VSX2+ retinal progenitors in day 35 retinal organoids [Scale: 40µm]. (C) Levels of 4/8 YAP1 isoforms in iPSCs, day 35 (D35) and day 60 (D60) retinal organoids. POM: periocular mesenchyme. pR: presumptive retina. pRPE: presumptive retinal pigmented epithelium. OF: optic fissure. iPSC: induced pluripotent stem cell.

### 3.2 Four YAP1 isoforms are expressed in human retinal organoids

We then studied YAP1 expression in VSX2+ retinal progenitor cells within early-stage hiPSC-derived ROs (day 35)(Fig. 1B). In these cells, YAP1 localisation was found to be predominantly cytoplasmic similar to the human embryo. At the RNA level, alternate splicing of YAP1 results in 8 mRNA isoforms. The main difference between YAP1 isoforms is the presence or absence of a second WW domain, with additional sequence differences within the transcriptional activation domain (TAD). Analysis of RNA-seq data from early-stage ROs (day 35 and day 60) revealed that four main YAP1 isoforms are being expressed: YAP1-201 (full length canonical isoform and most abundant, generating YAP1-2γ protein), YAP1-202 (WW2 domain absent, generating YAP1-1δ protein), YAP1-208 (shorter TAD, generating YAP1-2α protein), and YAP1-209 (additional sequence within TAD, generating YAP1-2β protein)(Fig. 1C).

### 3.3 YAP1 variant selection and in silico modelling

To gain insights into YAP1 structure and function, we studied five ClinVar-annotated YAP1 variants along with a previously unreported change (c.1068G>T, p.Glu356Asp) (Table 1). The latter was identified through clinical exome sequencing in a patient with coloboma, cleft palate, cleft of the alveolar ridge, ventricular septal defect and developmental delay (Table 1, Fig.2A). Clinical evaluation could not confirm the pathogenicity of the p.Glu356Asp variant, which was classified as a variant of uncertain significance (VUS). This classification was supported by its presence in the unaffected mother of the patient and in 13 alleles in the gnomAD v4 dataset. It is noted that the Glu356 residue is conserved across zebrafish, chicken, mouse and human, and that the variant is a conservative change from glutamate to another acidic side-chain aspartate.

**Figure 2.**
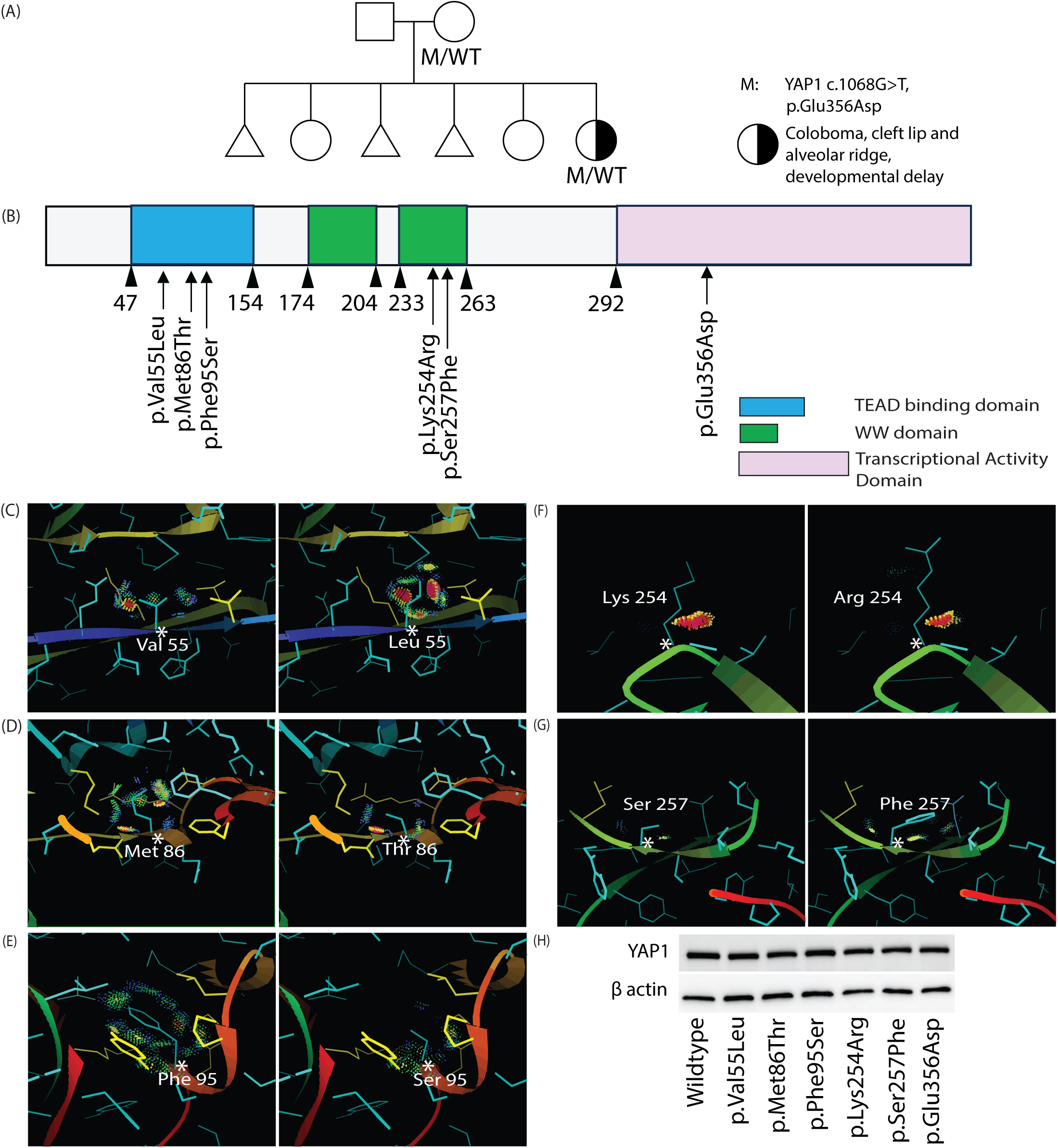
Modelling effects of YAP1 variants on protein structure. (A) Pedigree for YAP1 p.Glu356Asp. (B) Location of YAP1 variants from this study in different protein domains. (C-G) Introduction of YAP1 variants in solved protein structures. (C-E) YAP1-TEAD complex: p.Val55Leu, p.Met86Thr, p.Phe95Ser. (F-G) YAP1-LATS: p.Lys254Arg, p.Ser257Phe. Green and blue dots represent favourable van der Waals interactions, yellow and pink spikes represent unfavourable atomic overlaps. (H) Western blot showing non-degraded YAP1 variant proteins.

The studied variants are distributed across the domain structure of YAP1: p.Val55Leu, p.Met86Thr and p.Phe95Ser are located within the TEAD binding domain; p.Lys254Arg and p.Ser257Phe are found in the WW2 domain; and p.Glu356Asp is situated in the TAD (Fig. 2B). Using current variant effect predictors, pathogenicity scores varied across different algorithms. However, p.Phe95Ser and p.Ser257Phe were consistently predicted to be pathogenic (Table 2).

**Table 2:** Comparison of YAP1 pathogenicity and conservation prediction scores.

| Protein change | Pathogenicity scores |  |  |  |  | Conservation scores |  |  |
| --- | --- | --- | --- | --- | --- | --- | --- | --- |
|  | REVEL score | CADD | SIFT | Alpha Missense GRCh38 | PolyPhen2 | Mutation Taster | Mutation Assessor | PhyloP100 |
| p.Val55Leu | 0.035 | Quite likely deleterious | Tolerated | Pathogenic | Benign | Disease causing | Medium | 0.711 |
| p.Met86Thr | 0.141 | Quite likely deleterious | Damaging | Pathogenic | Benign | Disease causing | Medium | 2.122 |
| p.Phe95Ser | 0.189 | Probably deleterious | Damaging | Pathogenic | Probably damaging | Disease causing | Medium | 5.008 |
| p.Lys254Arg | 0.492 | Quite likely deleterious | Tolerated | Benign | Probably damaging | Disease causing | Neutral | 9.321 |
| p.Ser257Phe | 0.927 | Probably deleterious | Damaging | Pathogenic | Probably damaging | Disease causing | Medium | 7.902 |
| p.Glu356Asp | 0.158 | Potentially deleterious | Tolerated | Benign | Benign | Polymorphism | Neutral | 0.82 |

To assess how the six studied variants affect YAP1 structure and key binding interfaces, we modelled them in silico utilising two available YAP1 protein structures (Fig. 2C-G) [30, 31]. Notably, the YAP1-TEAD complex has three highly conserved interfaces (YAP1 residues 52-58, 61-73, and 86-100). The p.Val55Leu variant is found within the first interface (residues 52-58). This amino-acid substitution preserves hydrophobicity, but the larger side chain increases the number of interactions while also introducing van der Waals (vdW) clashes with neighbouring residues, potentially indicating a pathogenic effect. The third interface (residues 86-100) appears to be the most critical one; it is highly hydrophobic, with the side chains of Met86, Leu91, and Phe95 of YAP1 engaging a hydrophobic TEAD pocket and forming multiple vdW interactions [30]. A previous alanine scan of YAP1 residues found Met86, Arg89, Leu91, Ser94, Phe95 and Phe96 mutations resulted in a complete loss of activity. Moreover, increasing the hydrophobicity of Met86 and Phe95 enhanced YAP1-TEAD binding, highlighting their critical role within YAP1 [43]. From the protein modelling performed in this study, the p.Met86Thr and p.Phe95Ser variants are predicted to disrupt the hydrophobicity by switching to a polar residue, thus dampening the hydrophobic interface, reducing vdW interactions, creating a cavity at the site, and potentially decreasing YAP1-TEAD interactions.

With respect to YAP1 WW solved protein structure, WW domains of YAP1 interact with PPxY motifs of LATS [31]. YAP1 residues Lys254 and Asn250 Interact together to form the hydrophobic groove to fit the PPxY motif residue sidechains of LATS1. Thus, the p.Lys254Arg change is predicted to have an impact, although its functional relevance cannot be confirmed by modelling alone. For the p.Ser257Phe change, the wildtype serine participates in intra-molecular interactions that help form a β-sheet within YAP1. Substituting this small, polar residue with a bulky hydrophobic phenylalanine is likely to disrupt local interactions. Further, its placement on the protein surface may lead to reduced solubility due to unfavourable exposure of hydrophobic side chains. It is noted that structural modelling of the p.Glu356Asp variant was not feasible, as the affected residue is not present in any experimentally determined YAP1 structures and lies within a low-confidence region in AlphaFold models.

### 3.4 YAP1 variants do not cause protein degradation

To test if the YAP1 variant effects are due to protein degradation, the studied variants were introduced into a canonical full-length sequence plasmid (wildtype EYFP-YAP1) and were over-expressed in HEK293 LTV cells. Western blot showed similar expression levels of wildtype and mutant proteins (Fig. 2H), indicating that YAP1 variants do not induce protein degradation.

### 3.5 YAP1 variants p.Met86Thr and p.Ser257Phe differentially affect sub-cellular localisation

As noted above, organoid and human ocular tissue staining (Fig.1A-B) revealed YAP1 localisation in both the nucleus and cytoplasm, where distinct binding partners mediate compartment-specific functions. We hypothesised that missense changes in YAP1 may disrupt these interactions, including those with regulators of its subcellular localisation, such as LATS1/2. To assess the impact of the YAP1 variants on subcellular localisation, we generated stable HEK293 LTV cell lines expressing EYFP-tagged YAP1 constructs, including the phosphoacceptor site mutation p.Ser127Ala as positive control (known to impair phosphorylation, thus reducing cytoplasmic translocalisation) [44–46]. The EYFP intensity was measured in the nucleus and cytoplasm of the same cell for each variant to calculate the nuclear vs cytoplasmic localisation ratio. The YAP1 variants p.Val55Leu and p. Lys254Arg had no significant effect on YAP1 cellular localisation. Similarly, despite lying within the predicted nuclear export sequence (residues 340–357) [47], p.Glu356Asp had no effect. The TEAD binding domain variant p.Met86Thr showed a significant decrease in nuclear localisation, whereas p.Phe95Ser showed a non-significant decrease (Fig. 3A-B, Fig S1). Whereas the WW2 domain variant p.Ser257Phe significantly increased YAP1 nuclear localisation, the p.Lys254Arg variant did not (Fig. 3A-B, Fig S1). This indicates that Ser257 is key for regulating YAP1 localisation, likely through phosphorylation or LATS1/2 binding, consistent with protein modelling and the known interactions of both WW domains with LATS1/2 [31].

**Figure 3.**
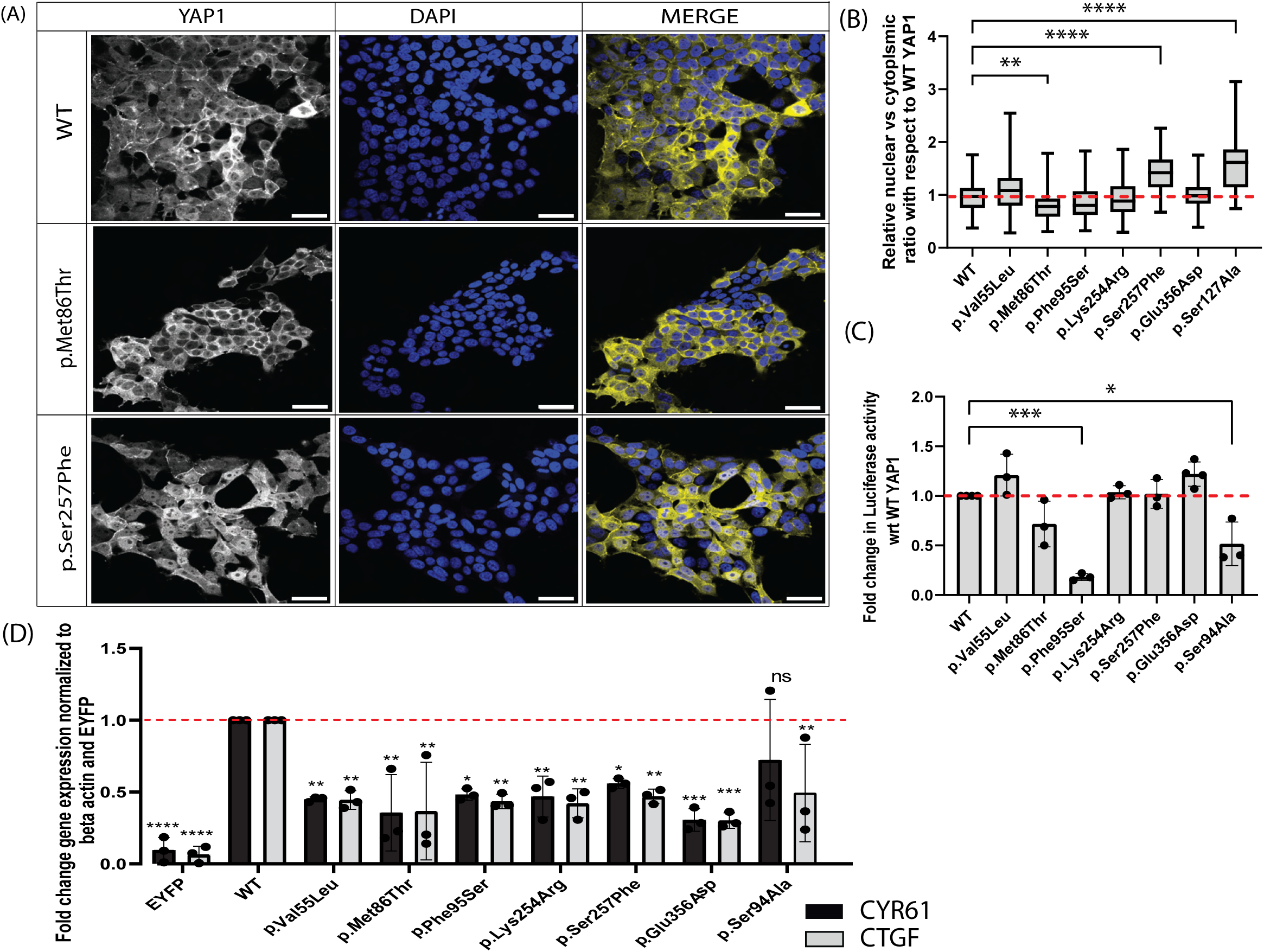
YAP1 variants affect protein localisation and transcriptional activity. (A) Sub-cellular localisation of EYFP-YAP1 wild-type (WT), p.Met86Thr and p.Ser257Phe variants. [Scale: 50µm] (B) Quantification of relative nuclear vs. cytoplasmic localisation of YAP1 variants with respect to WT EYFP-YAP1, with experimental control p.Ser127Ala. >100 cells analysed from 3 biological replicates (N=3, n≥100). Data presented as box and whiskers plot representing 25^th^ to 75^th^ percentile with a line at the median (C) Fold change in TEAD-dependent luciferase assay of EYFP-YAP1 WT, variants and control p.Ser94Ala. N=3. (D) Fold-change in expression of endogenous *CYR61* and *CTGF* by EYFP-YAP1 WT, variants and control p.Ser94Ala. N=3. Bar charts represent mean ± SD. Statistical significance was defined as: p < 0.05 (*), p < 0.01 (**), p < 0.001 (***), and p < 0.0001 (****); ns = not significant.

### 3.6 YAP1 variants in the TEAD binding domain affect transcriptional activity in luciferase assay

A key function of YAP1 is mediated through TEAD binding and activation of target genes. To assess if the studied YAP1 variants affect YAP1-TEAD dependent transcription, a luciferase assay with a synthetic TEAD-responsive promoter was performed by over-expression in HEK293 LTV cells. The positive control variant p.Ser94Ala (previously shown to reduce YAP1-TEAD interaction)[35, 48] reduced luciferase activity as expected. Out of the other TEAD binding domain variants, only p.Phe95Ser showed a significant reduction in transcriptional activity, although p.Met86Thr had a non-significant trend towards decreased activity (Fig. 3C). No effect on TEAD-dependent luciferase activity was observed for the p.Val55Leu, p.Lys254Arg and p.Glu356Asp variants (Fig. 3C). Although p.Ser257Phe had a higher nuclear localisation, it did not affect the transcriptional activity (Fig. 3B-C).

### 3.7 YAP1 variants in different domains downregulate endogenous downstream targets

To assess the effect of the studied YAP1 variants in a biologically-relevant chromatin environment, we compared the expression of endogenous *CTGF* and *CYR61* following over-expression of YAP1 variants in HEK293 LTV cells. Compared to the EYFP control vector, wildtype EYFP-YAP1 over-expression led to robust induction of *CTGF* and *CYR61* expression (Fig. 3D). All variants showed approximately a half-fold reduction in the induction of both genes, with the greatest reduction observed for p.Glu356Asp (Fig. 3D). Although only p.Phe95Ser significantly reduced TEAD-dependent luciferase activity, there is a consistent decrease in endogenous target gene induction across all variants. A possible explanation is that the TEAD-responsive reporter plasmid was episomal and highly accessible. In contrast, endogenous genes are subject to epigenetic regulation such as histone modifications and DNA methylation, which can restrict or enhance transcription factor access, potentially bypassing critical regulatory mechanisms. Further, endogenous genes may require cooperative binding of YAP1-TEAD to multiple factors, such as chromatin-remodelling complexes, or long-range enhancer interactions that are absent in the reporter [49]. This indicates that native gene context and chromatin environment are crucial for analysing the effect of missense variants on regulators of transcription.

### 3.8 YAP1 variants in the TEAD binding domain affect TEAD binding

To assess if the decrease in transcriptional activity observed from variants in the TEAD binding domain was due to decreased YAP1-TEAD binding, we measured YAP1-TEAD protein-protein interaction strength using FCCS. FCCS is a highly sensitive confocal microscopy technique that measures the simultaneous diffusion of two fluorescently labelled molecules to detect and quantify molecular interactions in live cells (in this case, cells over-expressing EYFP-YAP1 variants and TEAD1-mScarlet-I). By analysing the correlated fluctuations in EYFP and mScarlet-I fluorescence signals, FCCS provides quantitative estimates of YAP1-TEAD1 protein–protein binding affinity (Fig. 4A-C and Fig. S2). This showed that the wildtype YAP1-TEAD1 interaction had a mean dissociation constant (Kd) of 283 ± 72 nM (mean ± s.d), indicating moderate binding affinity (Fig. 4D and Fig. S2(C)). Among the studied variants, p.Met86Thr and p.Phe95Ser showed a significantly increased Kd (weaker force of interaction) similar to the control p.Ser94Ala (Fig. 4D). As would be expected for YAP1 that is not bound to TEAD or DNA, these variants also exhibited an increased rate of diffusion compared to wild-type (Fig. 4E). The YAP1-TEAD1 dissociation constant (inverse of binding affinity) negatively correlated with luciferase activity (Fig. 4F), suggesting that physical interaction between YAP1 and TEAD is necessary and rate-limiting for downstream gene activation in the luciferase assay.

**Figure 4.**
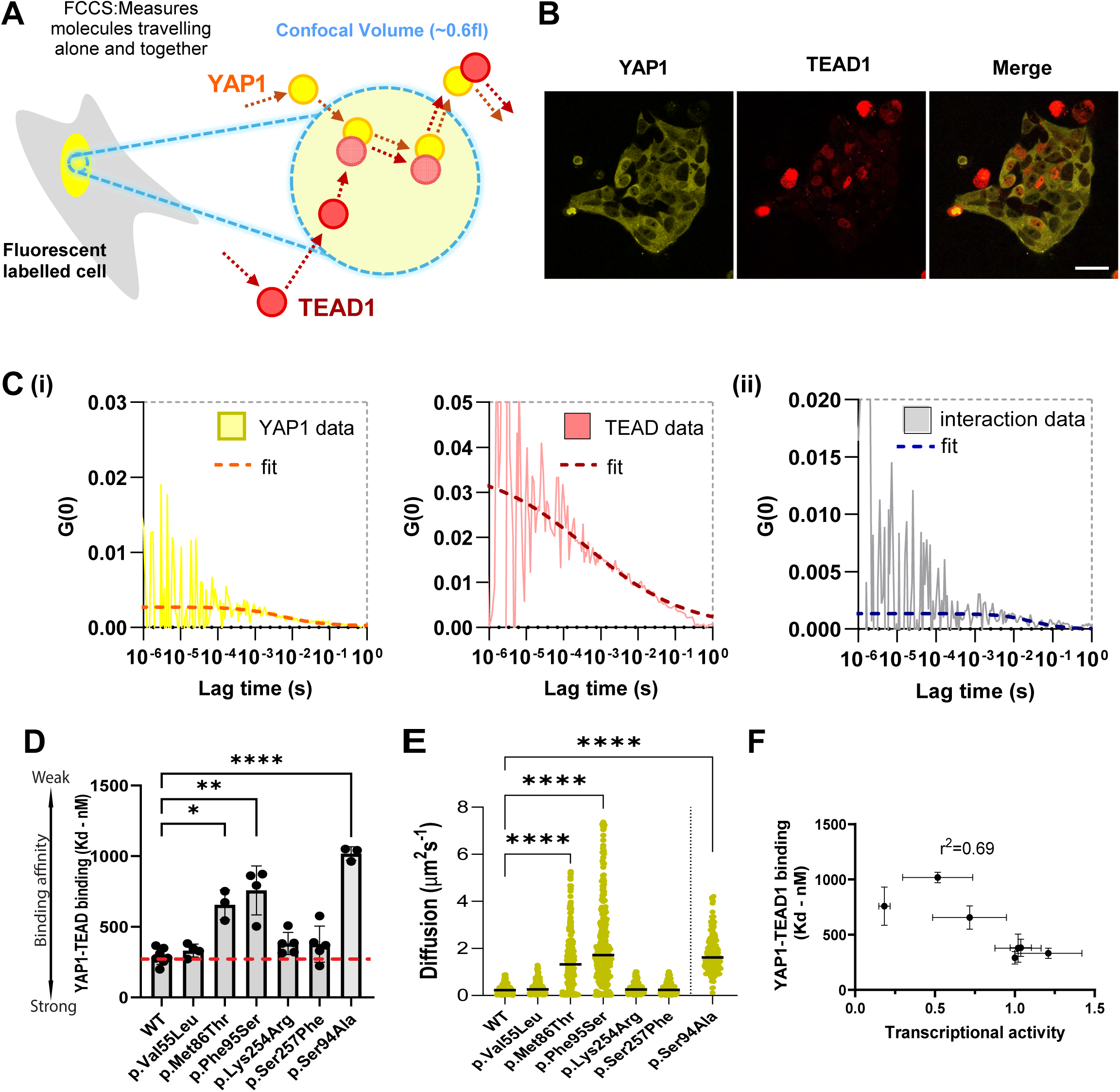
Quantification of YAP1-TEAD1 binding by fluorescence cross correlation spectroscopy (FCCS) (A) Schematic of FCCS principle to measure diffusion and protein-protein interaction strength of fluorescent tagged YAP1 and TEAD1 in the nucleus. (B) Representative image of YAP1 and TEAD1 expression in HEK293 LTV cells. (C) Representative (i) auto-and (ii) cross-correlation data showing raw data and fit lines for total EYFP-YAP1 and TEAD1-mScarlet and complexed fluorescent proteins (YAP1-TEAD1). (D) Dissociation constant (Kd) of YAP1-TEAD1 binding affinity of EYFP-YAP1 WT, variants and control p.Ser94Ala. N≥3. Bars show mean ± SD. (E) Diffusion rate for EYFP-YAP1 WT, variants and control p.Ser94Ala measured from auto-correlation data. All data points plotted with bars at mean. (F) Luciferase activity vs YAP1-TEAD1 dissociation constant (inverse of binding affinity) for EYFP-YAP1 WT and variants. Error bars are ± SD. Statistical significance was defined as: p < 0.05 (*), p < 0.01 (**), p < 0.001 (***), and p < 0.0001 (****); ns = not significant.

### 3.9 Inhibition of YAP1-TEAD binding in retinal organoids reduces proliferation and increases apoptosis

Functional testing of YAP1 variants in HEK293 LTV cells revealed that coloboma-associated mutations negatively affect YAP1–TEAD–dependent transcriptional activity, suggesting that disruption of nuclear YAP1 function may contribute to coloboma pathogenesis. However, staining of retinal tissue indicated YAP1 enriched in the cytoplasm. We next investigated whether YAP1 is also present in the nucleus (and undergoes active nucleo-cytoplasmic shuttling in retinal tissues) by treating early-stage ROs with the nuclear export inhibitor Selinexor (SLX). Staining for total YAP1 protein revealed a more pronounced YAP1 nuclear localisation in the SLX condition, as indicated by a significantly higher nuclear:cytoplasmic YAP1 ratio (Fig. 5A-B). The increase is consistent with SLX blocking XPO1-mediated nuclear export, preventing YAP1 from exiting the nucleus [50]. This suggests that YAP1 is dynamically shuttled between the nucleus and cytoplasm and that it potentially has a nuclear role regulating transcription in retinal progenitor cells.

**Figure 5.**
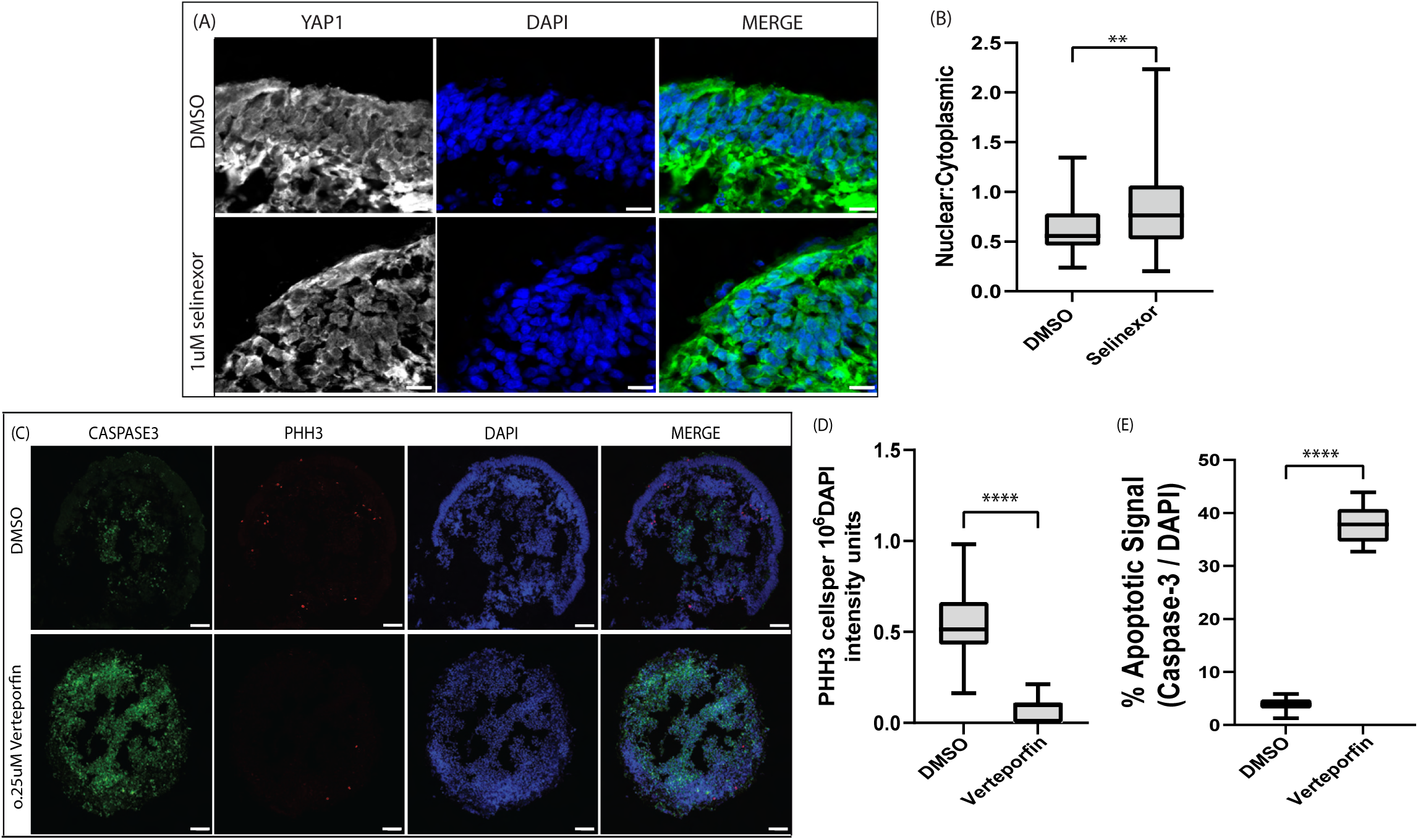
Functional role of YAP1 in day 40 retinal organoids. (A-B) YAP1 expression after 7 day selinexor (nuclear protein export inhibitor) treatment in organoids and quantification of YAP1 nucleus vs cytoplasmic localisation compared to DMSO control group. [Scale: 10µm]. (C) Cleaved caspase3 and Phosphohistone H3 (PHH3) expression in retinal organoids after 7 day o.25 uM Verteporfin (VPF; YAP1-TEAD inhibitor) treatment of organoids and control DMSO group [Scale: 50µm]. (D-E) Comparison of proliferative (PHH3+) and apoptotic (Caspase3+) cells after VPF treatment compared to DMSO control group. Quantification of PHH3+ cells per 10^6^ DAPI intensity units showed a significant reduction in proliferative activity in VPF-treated organoids compared to control. Apoptosis was calculated as the percentage of cleaved Caspase3 signal relative to total DAPI intensity per region of interest (ROI). Data is presented as mean ± SD. N=3. Statistical significance was defined as: p < 0.05 (*), p < 0.01 (**), p < 0.001 (***), and p < 0.0001 (****); ns = not significant.

To test YAP1 transcriptional function, early-stage ROs were treated with verteporfin, an inhibitor of YAP1-TEAD binding, for seven days [51]. We then evaluated retinal progenitor proliferation and apoptosis by assessing the number of phospho-histone H3 and cleaved caspase-3 positive cells, respectively. After 0.25uM verteporfin treatment, we observed a significant decrease in proliferating cells and a significant increase in cells undergoing apoptosis (Fig. 5C-E). This indicates a key role of YAP1-TEAD dependent transcription in human retinal progenitor proliferation and maintenance during early retinal development.

## 4. DISCUSSION

Determining the pathogenicity of YAP1 missense variants associated with coloboma requires integrating evidence from gene expression studies, mechanistic knowledge of gene function, and case-level data. Here, we show that YAP1 is expressed across multiple tissues in the developing eye surrounding the optic fissure, including the periocular mesenchyme, neural retina and RPE. Transcriptomic analysis of early-stage ROs revealed four main YAP1 splice isoforms, three of which contain two WW domains and would be affected by variants in the second WW domain. An alternative TSS isoform (Met179; ENSMBL ID: 203), previously proposed to bypass nonsense-mediated decay and partially rescue YAP1 haploinsufficiency, was not detected [4]. Even if expressed, this isoform would lack the TEAD-binding domain, limiting its ability to restore full YAP1 function.

In silico prediction tools like REVEL and AlphaMissense offer powerful approaches for predicting the potential impact of missense variants on protein function [23, 26]. However, out of the six YAP1 variants tested here, only two, p.Phe95Ser and p.Ser257Phe were consistently predicted as pathogenic across multiple in silico tools. Structural modelling further supported the pathogenicity of these variants. In addition, our modelling suggested that p.Val55Leu is likely pathogenic due to vdW clashes, while p.Met86Thr may be pathogenic due to the introduction of a polar side chain into the key hydrophobic TEAD-binding interface. Consistent with the predictions, our in vitro studies revealed alterations in localisation, transcriptional activity, or TEAD binding for these variants. Notably, some variants consistently predicted to be benign by computational tools still resulted in reduced induction of endogenous YAP1–TEAD target genes, highlighting the limitations of current in silico approaches and the value of functional validation. Unsurprisingly, our in vitro functional testing of YAP1 revealed assay-dependent effects. For example, TEAD-binding domain variants had the largest effect in a TEAD-dependent luciferase assay, whereas a WW domain variant primarily affected YAP1 sub-cellular localisation. These findings likely reflect the distinct interactions of YAP1 domains with different protein partners, suggesting that variants in separate domains drive pathogenicity through different mechanisms (Table 3).

**Table 3:**
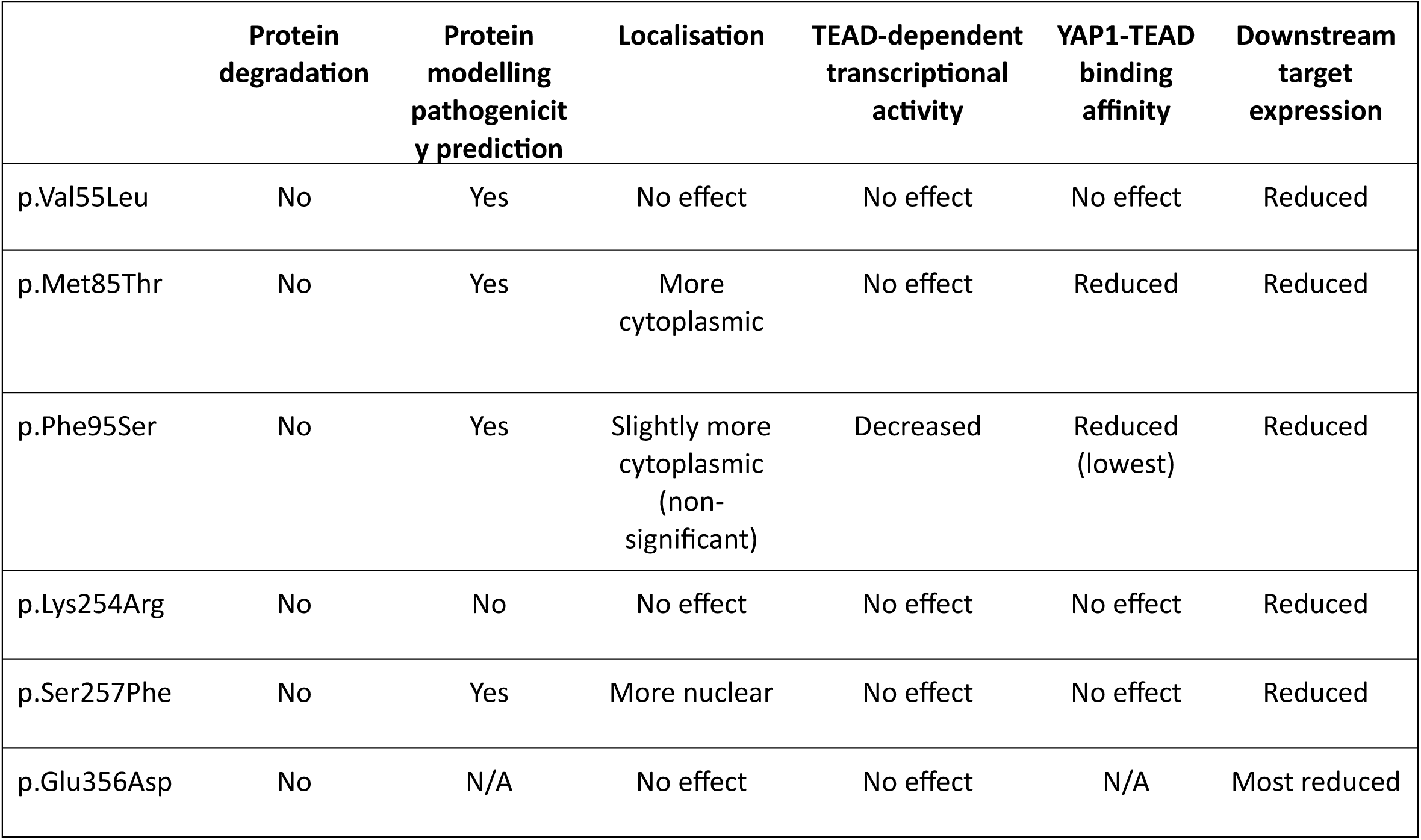
Comparison of experimental results from YAP1 variant assays.

A key similarity was that all variants were significantly less effective at inducing the expression of endogenous target genes *CYR61* and *CTGF* compared to wildtype YAP1, with the TAD variant (p.Glu356Asp) having the most pronounced effect. The TAD mediates interactions with chromatin-modifying complexes and other proteins that regulate the transcription machinery, suggesting that the studied TAD variant would affect all YAP1-dependent transcription [52].

To assess the impact of disrupted YAP1 transcription in a relevant context, early-stage ROs were treated with verteporfin, a YAP1-TEAD interaction inhibitor. This revealed that the YAP1-TEAD interaction is a positive regulator of retinal progenitor cell proliferation and is important for human retinal development. This is consistent with observations from a retinal-specific YAP1 knockout mouse model, which exhibited reduced retinal progenitor proliferation, increased apoptosis, coloboma and microphthalmia [8]. Importantly in human studies, TEAD binding sites were found to be enriched in retinal progenitor cells in organoids by ATAC-seq [53]. Also, treatment of organoids with MGH-CP1, a TEAD inhibitor, resulted in: a decrease in the retinal progenitor marker VSX2; a decrease in proliferating cells; and a small increase in apoptosis [53]. These observations suggest that, in addition to YAP1, missense variants in TEAD transcription factors may also lead to ocular disorders such as coloboma. Interestingly, a variant in TEAD1 p.Tyr421His, has been reported to cause a chorioretinopathy with coloboma-like defects in the retina, RPE and choroid near the optic nerve [54, 55].

Our findings suggest that YAP1–TEAD transcriptional activity is essential for maintaining proliferative retinal progenitors, and that reduced proliferation in patients with YAP1 variants may disrupt retinal morphology and hinder optic fissure closure. However, there are likely to be other YAP1-dependent processes important for optic fissure closure that are missing from the retinal organoid model. Prior to fusion in the embryo, the cells of the optic fissure margin undergo basement membrane degradation, apical remodelling, EMT-like transitions and subsequent cell mixing and reorganisation to form continuous cell layers of neural retina and RPE [56, 57]. YAP1 mechanosensing at the fissure margins may regulate many of these dynamic processes, likely in combination with other pathways necessary for ocular development. For example, TGF-β signalling, a key regulator of ECM remodelling is essential for this process and TGF-β2 inactivation in mouse led to a failure in optic fissure fusion and coloboma [58]. Both YAP1-1 and YAP1-2 mediate EMT, but TGF-β dominantly increases YAP1-2 stability, enhances its nuclear localisation, and promotes cell migration, [59]. Variants in the second WW2 domain could impair this TGF-β dependent EMT of YAP1-2, thereby contributing to coloboma. Similarly, precise regulation of BMP signals is also important for optic fissure closure. YAP1 can bind via it’s two WW domains to SMAD1/5, the transcriptional mediators of the BMP pathway, and potentially influence downstream targets [60]. Furthermore, abrogation of the Wnt pathway and β-catenin leads to coloboma and increased cytoplasmic YAP1 enhances recruitment of β-catenin to its destruction complex, thereby reducing WNT target gene expression and affecting RPE differentiation [61–63].

In conclusion, this study provides new insights into YAP1 function during early human retinal development by combining organoid studies with functional assessment of disease-associated variants. This combined approach enables a deeper understanding of YAP1’s role in human tissue development and offers a foundation for interpreting the clinical significance of rare YAP1 variants.

## Ethical Approval

Human embryonic material was collected from medical and surgical terminations of pregnancy under ethical approval from the North West Research Ethics Committee (23/NW/0039) and under the codes of practice issued by the Human Tissue Authority and legislation of the UK Human Tissue Act 2008. Informed consent was obtained from all participants. For the data from coloboma proband, the Informed consent for inclusion in the research study was signed by the family. The consent form was approved by the Self Regional Hospital institutional review board.

## Acknowledgments

We thank members of the Manning lab for their critical feedback and continuing support. The authors also thank the Bioimaging Facility of the University of Manchester for technical support.

## Author Contribution Statement

Conceptualization: C.M., P.S., F.M., S.S., A.R-L., G.A.; Methodology: C.M., P.S., S.S., A.R-L., A.M., F.M., J.B., R.J., L.B.; Investigation: S.S, A.R-L., M.S., X.Y., R.L., L.B., J.M.D., J.R.J., E.S.; Formal analysis: S.S; A.R-L., M.S., J.B., S.L.; Writing -original draft: S.S., C.M., Writing -review & editing: S.S., C.M, P.S.; Visualisation: S.S; A.R-L., J.B., S.L.; Supervision: C.M., P.S., F.M., J.E., A.M., G.A.

