## Supplemental figures 1-2 for "Domain-specific mechanisms of YAP1 variants in ocular coloboma revealed by in-vitro and organoid studies": supp figure 1-2_combined.pdf

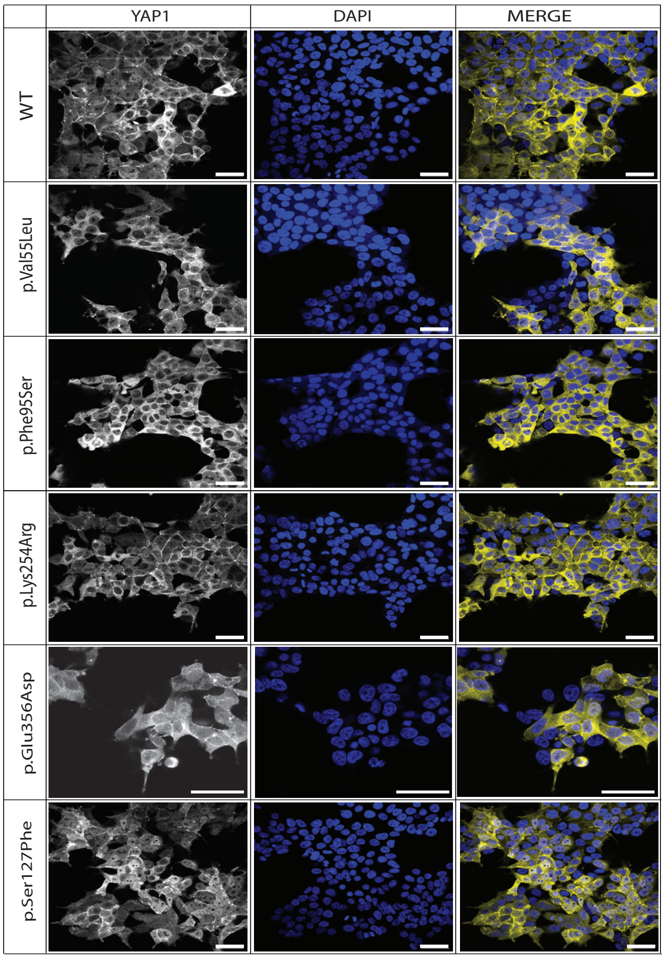


Supplementary figure 1. YAP1 sub-cellular localisation.
Representative images of HEK293 LTV cells stably expressing EYFP-YAP1 WT and variants.
Blue – DAPI. Yellow – EYFP-YAP1. [Scale: 50μm]

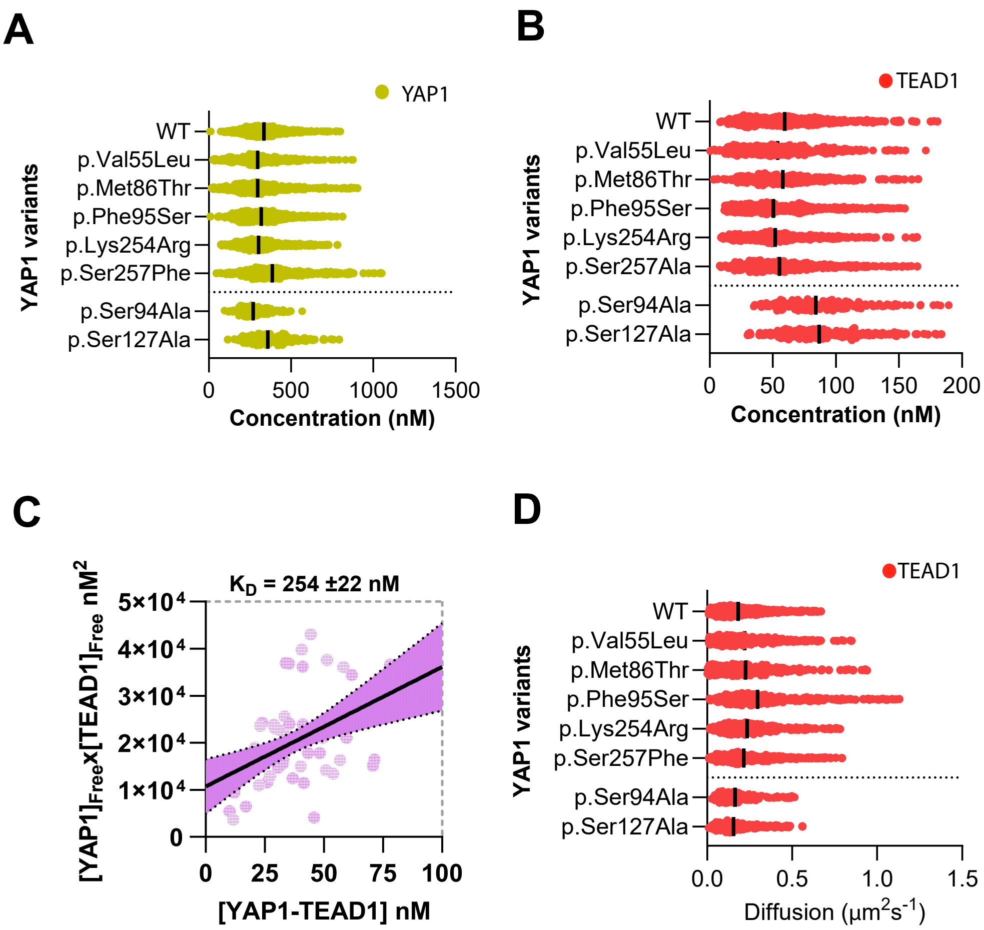


Supplementary figure 2. Quantification of YAP1-TEAD1 binding by fluorescence cross correlation spectroscopy (FCCS). Protein concentration of (A) YAP1 and (B) TEAD1 respectively from auto-correlation data. All data points plotted with bars at median. (C) Representative dissociation plot to determine KD for EYFP-YAP1 WT and TEAD1-mScarlet from FCCS data. (D) TEAD1 diffusion rate when co-expressed with EYFP-YAP1 WT and variants from auto-correlation data. All data points plotted with bars at median.
